# Hurricane-induced structural disturbance amplifies forest vulnerability to subsequent heatwaves

**DOI:** 10.64898/2026.08.10.743363

**Authors:** Jingyu Dai, Anna B. Harper, Xinyou Li, Gabriel J. Kooperman, Maria Uriarte, Thomas L. Mote

## Abstract

Sequential hurricane-heatwave events threaten forest resilience via understudied legacy effects. Using Bayesian Structural Time Series, piecewise Structural Equation Modeling, and a 21-event global synthesis, we quantify how structural degradation non-linearly amplifies productivity loss during subsequent heatwaves. Our Hurricane Michael (2018) case study reveals significant negative GPP legacy effects during the 2019 heatwave. Intact, tall and diverse canopies buffer microclimates and moderate thermal sensitivity. Hurricane-induced structural simplification removes this protection, exposing temperature-sensitive shaded leaves to extreme stress. We identified a context-dependent hydraulic trade-off: structural complexity provides shading but exacerbates forest sensitivity to water deficits during peak heat, the vulnerability of which reverses during the recovery phase. Globally, these legacy effects are triggered by heatwave intensity and modulated by soil type, with loamy-soil forests most vulnerable. These findings highlight the critical role of forest structure in forest responses to compound disturbances. Neglecting structural legacies in Earth System Models likely underestimates risks to global carbon sinks.

## Introduction

Climate change has increased the global frequency of hurricanes, heatwaves, and droughts, heightening the probability of sequential or simultaneous disturbances^1^. While concurrent events are well-studied (often with intervals ≤1 year)^2, 3^, sequential compound events pose a unique but understudied threat. An initial disturbance can erode ecosystem resilience, leaving a "legacy effect" that amplifies the impact of a subsequent, seemingly unrelated stressor^4, 5^. The legacy effect of a preceding event, such as physical damage or physiological stress, can predispose forests to a state of heightened sensitivity, potentially leading to non-linear impacts to forest dynamics and carbon flux when a second event strikes^5, 6, 7^.

Among these sequential compound events, the hurricane-heatwave disturbance represents a critical yet understudied biophysical mechanism. The capacity of a forest to withstand and recover from disturbance is fundamentally mediated by its structural state. Canopy height, structural and species diversity, and vertical complexity determine how effectively a forest buffers its internal microclimate and moderates the sensitivity of carbon fluxes to external stressors. Hurricanes act as profound structural disruptors, instantaneously reducing canopy height and diversity through defoliation, branch snapping, and tree mortality^8, 9^. By dismantling the structural foundations of microclimate regulation, hurricanes may therefore leave forests in a state of heightened sensitivity to subsequent climate extremes - a latent vulnerability that remains largely unquantified^9^. Forest structure, particularly canopy height, is key to microclimate buffering: taller canopies regulate localized land surface temperature (LST, ℃) and vapor pressure deficit (VPD, kPa) through shading, air mixing and evapotranspirative cooling^10, 11, 12^. Meanwhile, structural diversity indicates niche diversity^13^, such as the difference between shaded and sunlit leaves^14^, which enables varied and asynchronous individual responses to rising temperature and water deficit, thus moderating forest sensitivity to climate dynamics^15, 16^. We hypothesize that hurricane-induced structural simplification compromises the buffering and moderating capacity of an undisturbed forest. Consequently, when a heatwave follows, the disturbed forest may experience a more extreme microclimate and higher sensitivity than an intact forest would, leading to amplified suppression of forest gross primary productivity (GPP).

The catastrophic Hurricane Michael (Category 5, 2018) provides a unique "natural experiment" to test this hypothesis. It caused extensive forest structural disturbance across the southeastern U.S.^17^, transforming local longleaf pine woodlands from net carbon sinks into net sources^18^. The hurricane was followed by a record-breaking heatwave and drought in September 2019^19^. This sequence offers an ideal window to disentangle the cascading effects of structural damage on forest heat vulnerability. Here, we integrate multi-source remote sensing data with a statistically-informed counterfactual approach to quantify the impact of hurricane-heatwave disturbances on forest productivity. We first conduct a detailed mechanistic study on Hurricane Michael, and subsequently extend our analysis to a global scale to examine if these patterns are modulated by varying stand conditions. Specifically, we address three questions:

1. Did Hurricane Michael cause larger negative GPP anomalies in forests during the subsequent heatwave compared to the counterfactual GPP scenario?
2. How did forest structural traits and vertical canopy positions mediate this process during the heatwave peak and recovery phases?
3. Is the non-linear amplification in forest heatwave vulnerability consistent across global hurricane-heatwave compound events?

## Results

### Immediate impact and legacy effect of Hurricane Michael on ΔGPP

To capture the immediate physiological and structural damage caused by Hurricane Michael, we focused on the minimum ΔGPP (most negative GPP anomaly relative to a multi-year baseline) within the first three months post-landfall (October-December, 2018, **Figure 1a**) in hurricane-disturbed and undisturbed forests (**Methods**). During this three-month window, the hurricane-disturbed forests exhibited a pronounced and widespread decline in productivity. Mean ΔGPP loss across the disturbed pixels was 28.96 gC m^-2^ month^-1^, significantly higher than the reference forests (10.65 gC m^-2^ month^-1^). The most severe impacts were concentrated in coastal regions near the hurricane’s landfall point, where localized ΔGPP losses reached as high as 77.30 gC m^-2^ month^-1^. These results confirm that our sampling strategy effectively distinguished between forests experiencing wind damage and those remaining within the range of normal environmental variability.

**Figure 1.**
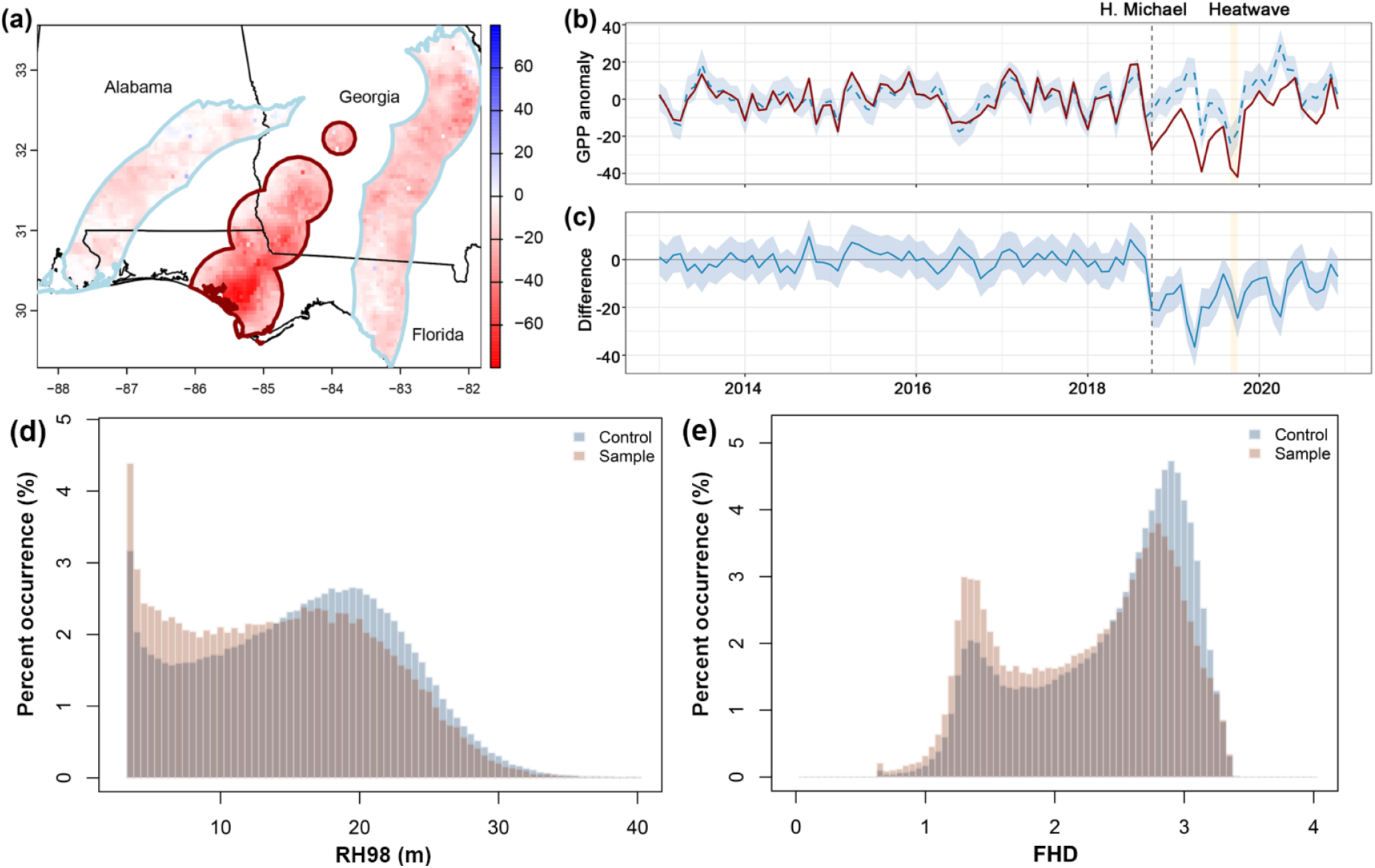
(a) Spatial distribution of the minimum GPP anomaly within the first three months following Hurricane Michael (Oct.-Dec. 2018). The hurricane-disturbed forests and the reference group are delineated by dark red and light blue boundaries, respectively. Red pixels indicate GPP losses relative to the pre-hurricane baseline. **(b)** Comparison of observed and counterfactual GPP anomaly time series for the hurricane-disturbed region. The blue dashed line represents the counterfactual GPP anomaly predicted by the BSTS model, with the light blue shaded area indicating the 95% credible interval. The dark red solid line represents the observed GPP anomaly time series. **(c)** The pointwise causal impact, calculated as the difference between the observed and predicted GPP anomalies, with the shaded area representing the 95% confidence interval. In panels **(b, c)**, the black vertical dashed line indicates the landfall of Hurricane Michael (Oct. 2018), and the orange rectangular shading highlights the period of the subsequent extreme heatwave (Sept. 2019). **(d-e)** Histograms illustrate the percentage distributions of canopy height (RH98) and Foliage Height Diversity (FHD). Red distributions represent the hurricane-disturbed forests, while blue distributions denote the undisturbed reference group.

Using a Bayesian Structural Time Series (BSTS) model trained on undisturbed reference forests to generate counterfactual ΔGPP trajectories (**Methods**), we demonstrated high predictive accuracy during the five-year pre-hurricane training period (September 2013 - September 2018), with no significant divergence observed between the actual ΔGPP observations and the model-predicted counterfactuals (**Figure 1b-c**). Following the landfall of Hurricane Michael in October 2018, a sustained and significant suppression of ΔGPP was observed. From October 2018 to December 2020, the observed ΔGPP remained consistently lower than the counterfactual predictions. While the magnitude of this divergence exhibited a gradual, fluctuating decline, which indicates a trajectory of recovery, the forest productivity had not fully returned to its expected baseline by the end of 2020. On average, the hurricane-disturbed forests experienced an extra ΔGPP loss of 14.24 gC m^-2^ month^-1^ throughout the post-landfall duration (95% CI = [12.16, 16.22]). During the heatwave in September 2019, the gap between observed and counterfactual ΔGPP widened sharply. The average extra GPP loss intensified to 18.76 gC m^-2^ month^-1^ (95% CI = [10.71, 26.83]) during the heatwave (**Figure 1b-c**). This amplification suggests that the forest degradation caused by Hurricane Michael significantly increased the forest’s vulnerability to the following heatwave, leading to an exaggerated decline in productivity.

## Hurricane Michael impacted forest structure and forest heatwave vulnerability

We characterized the structural impact of Hurricane Michael by quantifying the divergence in canopy structure between the hurricane-disturbed sample forests and the undisturbed reference group. The frequency distributions (**Figure 1d-e**) revealed a significant downward shift in both canopy height (indicated by the relative height metrics at 98%, RH98) and structure diversity (indicated by the foliar height diversity index, FHD, **Methods**) within the disturbed regions. Compared to the reference group, the sample forests exhibited a markedly lower proportion of pixels characterized by mature, tall canopies (RH98 ≈ 20m) and high structural diversity (FHD ≥ 3.0). Conversely, there was a disproportionate increase in the frequency of pixels with low canopy heights (RH98 < 5 m) and simplified vertical profiles (FHD ∈ [1.0, 2.0]). These distributional shifts indicate that the intense wind forcing of Hurricane Michael triggered widespread structural degradation, effectively converting a substantial portion of mature, diverse-structured forests into shorter, simplified woodlands.

The influence of forest structure on the microclimate and the sensitivity of ΔGPP to environmental stress showed a distinct reversal between the heatwave peak (September 2019) and the subsequent recovery phase (October 2019, **Figure 2**). In September 2019, the sampling region experienced severe thermal and hydrological stress, with mean ΔLST reaching 4.47 ± 1.02 ℃ and mean ΔVPD at 0.82 ± 0.09 kPa. We observed significant negative correlations between RH98 and both climate anomalies (slopes: -0.017 for ΔLST and -9.73×10^-4^ for ΔVPD; both p < 0.0001, **Figure 2a-b**), suggesting a clear buffering effect of taller forests against extreme heat and water stress. This negative relationship was most apparent in pixels with RH98 < 20 m, whereas in taller forests > 20 m, the buffering capacity appeared to saturate or weaken. Grouped regression analyses further revealed a dual moderating role of canopy height on GPP sensitivity. Taller forests exhibited less negative ΔLST-ΔGPP slopes but more negative ΔVPD- ΔGPP slopes (**Figure 2c-d**). These findings were validated by the piecewise Structural Equation Model (pSEM, **Methods**), where RH98 showed significant negative paths to ΔLST (standardized estimate: -0.11) and ΔVPD (-0.04). While both ΔLST and ΔVPD strongly suppressed ΔGPP (- 0.39 and -0.13, respectively), the positive interaction between RH98 and ΔLST (+0.04) and the negative interaction with ΔVPD (-0.07) confirm that forest structure moderates the sensitivity of productivity to climate stressors: taller structures buffer the negative impact of ΔLST on ΔGPP, but exacerbate the negative effect of ΔVPD on ΔGPP, during peak heatwave (**Figure 2e**).

**Figure 2.**
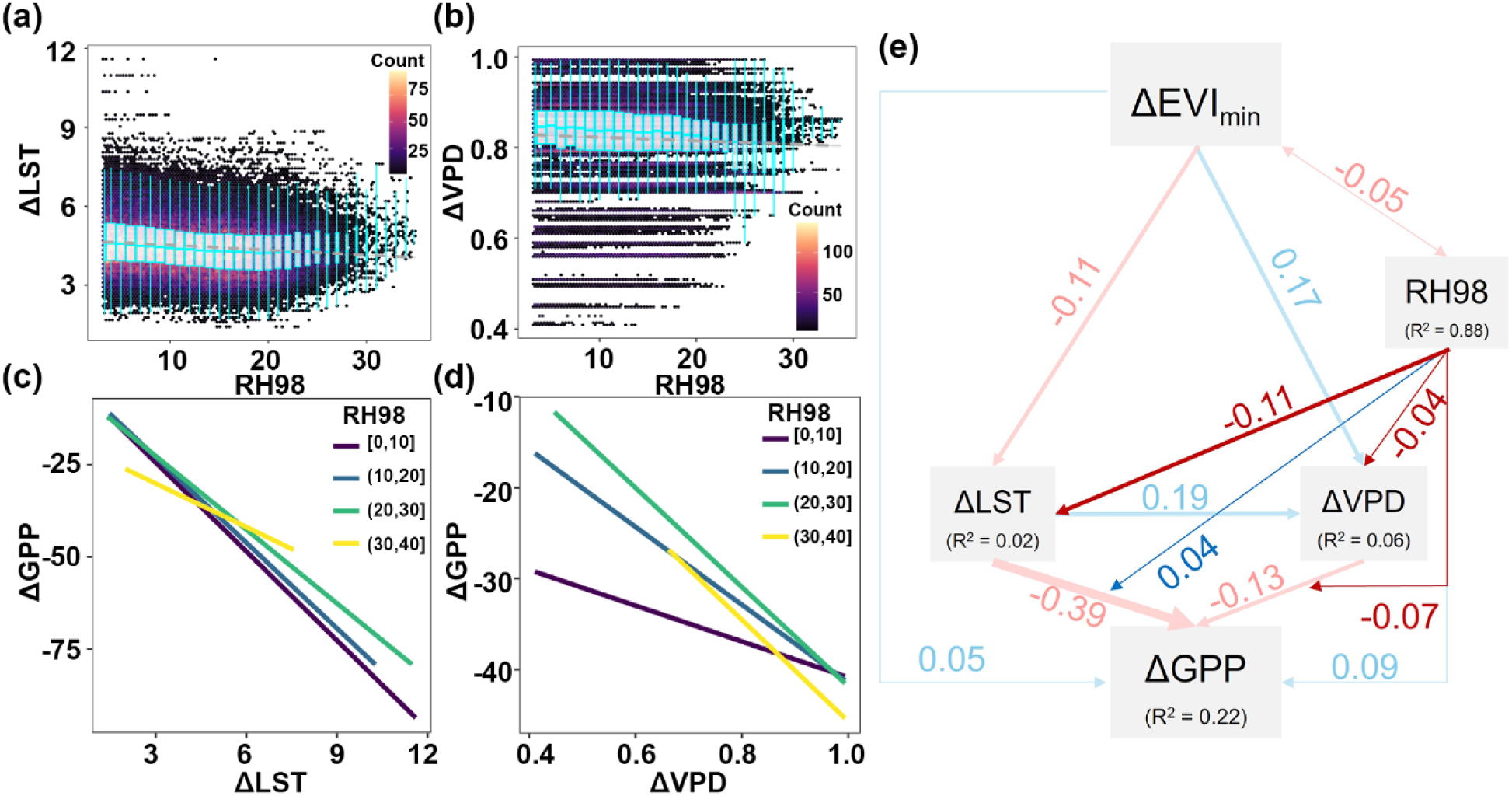
(a, **b)** Relationships between canopy height (RH98) and anomalies in Land Surface Temperature (ΔLST) and Vapor Pressure Deficit (ΔVPD) during the heatwave peak (Sept. 2019). The background density plots show the distribution of the raw pixel-level data. Boxplots represent the trends across 1-meter canopy height bins, while the dashed gray lines show the linear regression. **(c, d)** Grouped regressions between ΔGPP and ΔLST or ΔVPD indicating the sensitivity of ΔGPP to environmental stress as a function of forest height. More negative regression slopes indicate a higher vulnerability of ΔGPP to environmental fluctuations. **(e)** Piecewise Structural Equation Model (pSEM) illustrating the regulatory network between hurricane legacy effects (ΔEVImin), RH98 and environmental factors during the heatwave peak. Arrows represent significant causal paths (p ≤ 0.05), with blue/light-blue and red/light-red indicating positive and negative effects, respectively (thinner lines denote weaker associations). Highlighted paths (red/blue arrows) emphasize the buffering and moderating effects of RH98.

In October 2019, after the heatwave peak, environmental anomalies returned toward baseline levels (ΔLST = 0.75 ±1.30℃; ΔVPD = 0.01 ±0.05 kPa). During this recovery phase, the relationship between forest structure and microclimate reversed, with RH98 exhibiting positive regression slopes with both ΔLST (slope = 0.003, p < 0.0001, **Figure 3a**) and ΔVPD (slope = 0.001, p < 0.0001, **Figure 3b**). The moderating effect of forest structure also shifted. Grouped regressions indicated that taller forests’ ΔGPP possessed less sensitivity to both thermal and hydrological anomalies, showing less negative slopes for both ΔLST and ΔVPD (**Figure 3c-d**). The pSEM supported this transition (**Figure 3e**): the buffering effect of RH98 on ΔLST became non-significant (p > 0.05), while a positive path emerged between RH98 and ΔVPD (+0.10). While the impact of microclimates on ΔGPP are negative (ΔLST-ΔGPP: -0.26, ΔVPD-ΔGPP: - 0.24), the interaction terms between RH98 and both climate stressors were significantly positive (+0.04 for ΔLST and +0.14 for ΔVPD). These results collectively demonstrate that during the recovery period, taller canopies effectively lowered the sensitivity of ΔGPP to residual environmental fluctuations, facilitating a more robust post-heatwave recovery compared to shorter forest stands.

**Figure 3.**
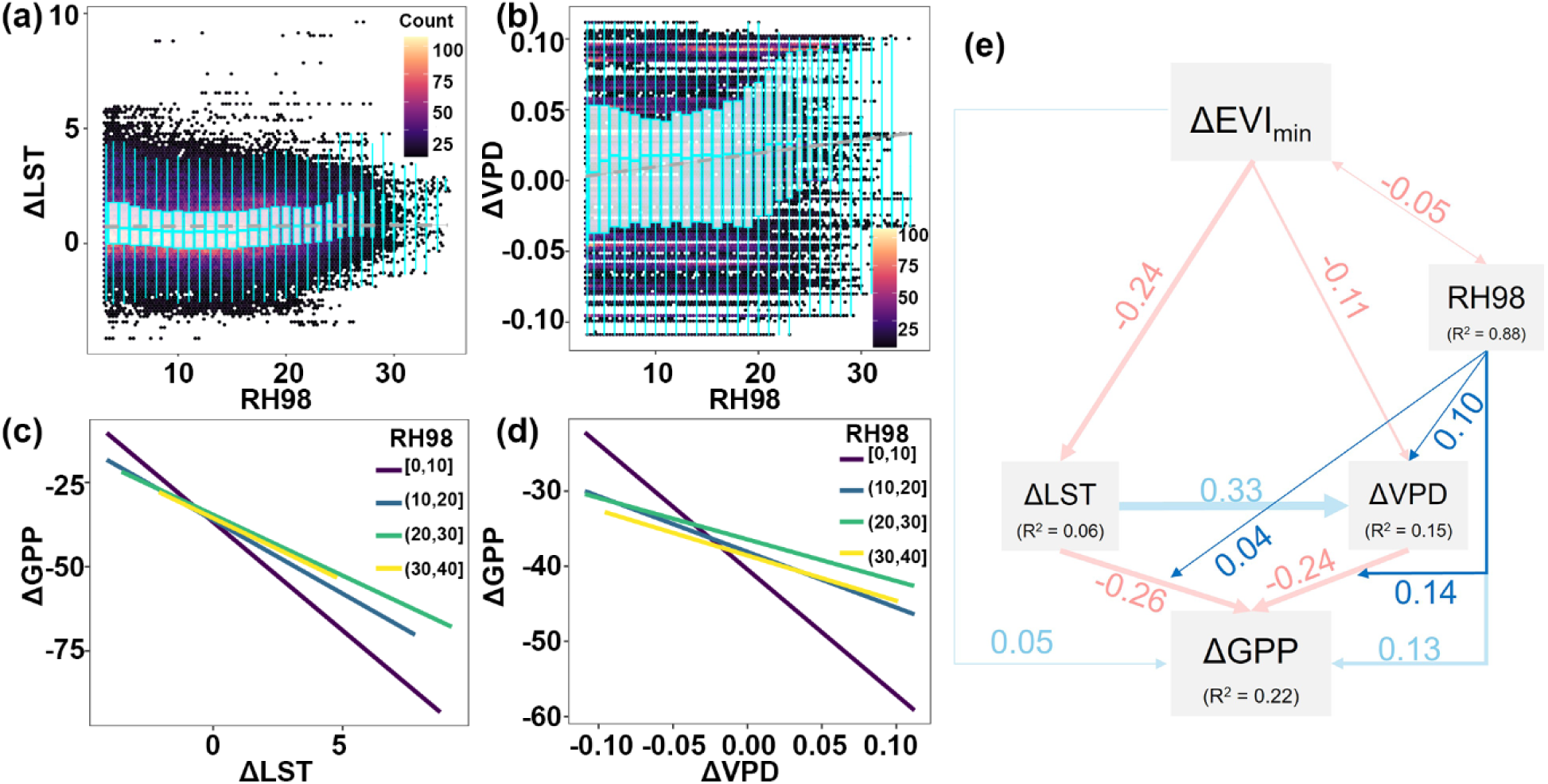
(a, **b)** Relationships between canopy height (RH98) and anomalies in Land Surface Temperature (ΔLST) and Vapor Pressure Deficit (ΔVPD) during recovery period (Oct. 2019). The background density plots show the distribution of the raw pixel-level data. Boxplots represent the trends across 1-meter canopy height bins, while the dashed gray lines show the linear regression. **(c, d)** Grouped regressions between ΔGPP and ΔLST or ΔVPD indicating the sensitivity of ΔGPP to environmental stress as a function of forest height. More negative regression slopes indicate a higher vulnerability of ΔGPP to environmental fluctuations. **(e)** Piecewise Structural Equation Model (pSEM) illustrating the regulatory network between hurricane legacy effects (ΔEVImin), RH98 and environmental factors during the heatwave peak. Arrows represent significant causal paths (p ≤ 0.05), with blue/light-blue and red/light-red indicating positive and negative effects, respectively (thinner lines denote weaker associations). Highlighted paths (red/blue arrows) emphasize the buffering and moderating effects of RH98.

To further elucidate the vertical heterogeneity of forest physiological responses, we conducted partitioned pSEM analyses for shaded (ΔGPPshaded) and sunlit GPP (ΔGPPsunlit) during the peak heatwave and recovery periods (**Figure 4**). Overall, the regulatory pathways involving RH98, microclimate, and the partitioned GPP components largely mirrored the patterns observed for total GPP in their respective months. During the peak heatwave (September 2019), the positive interaction between RH98 and ΔLST mitigated the sensitivity of both ΔGPPshaded and ΔGPPsunlit to thermal stress (+0.05 and +0.01, respectively), while the negative interaction between RH98 and ΔVPD amplified their vulnerability to water deficits (- 0.08 and -0.04, respectively). During the recovery period (October 2019), the interaction between RH98 and ΔVPD transitioned to positive (+0.12 and +0.13, respectively), reducing the sensitivity of both components to hydrological anomalies. Distinctive physiological sensitivities emerged when comparing relative importance of ΔLST and ΔVPD. In both months, ΔGPPshaded exhibited a markedly higher sensitivity to ΔLST (-0.45 and -0.32, respectively) than to ΔVPD (- 0.06 and -0.27, respectively), as evidenced by the larger absolute values of standardized path coefficients. In contrast, ΔGPPsunlit was more sensitive to ΔVPD (-0.23 and -0.11, respectively) than to ΔLST (-0.18 and -0.10, respectively). Furthermore, the pSEM consistently yielded higher explanatory power for ΔGPPshaded (R^2^ = 0.26 for Sept.; R^2^ = 0.31 for Oct.) compared to ΔGPPsunlit (R^2^ = 0.12 for Sept.; R^2^ = 0.06 for Oct.), which suggests that the regulation of forest structure and local microclimate are more profoundly exerted on the shaded sub-canopy layers, whereas sunlit leaves may be subject to large-scale meteorological drivers not fully captured by the local structural model.

**Figure 4.**
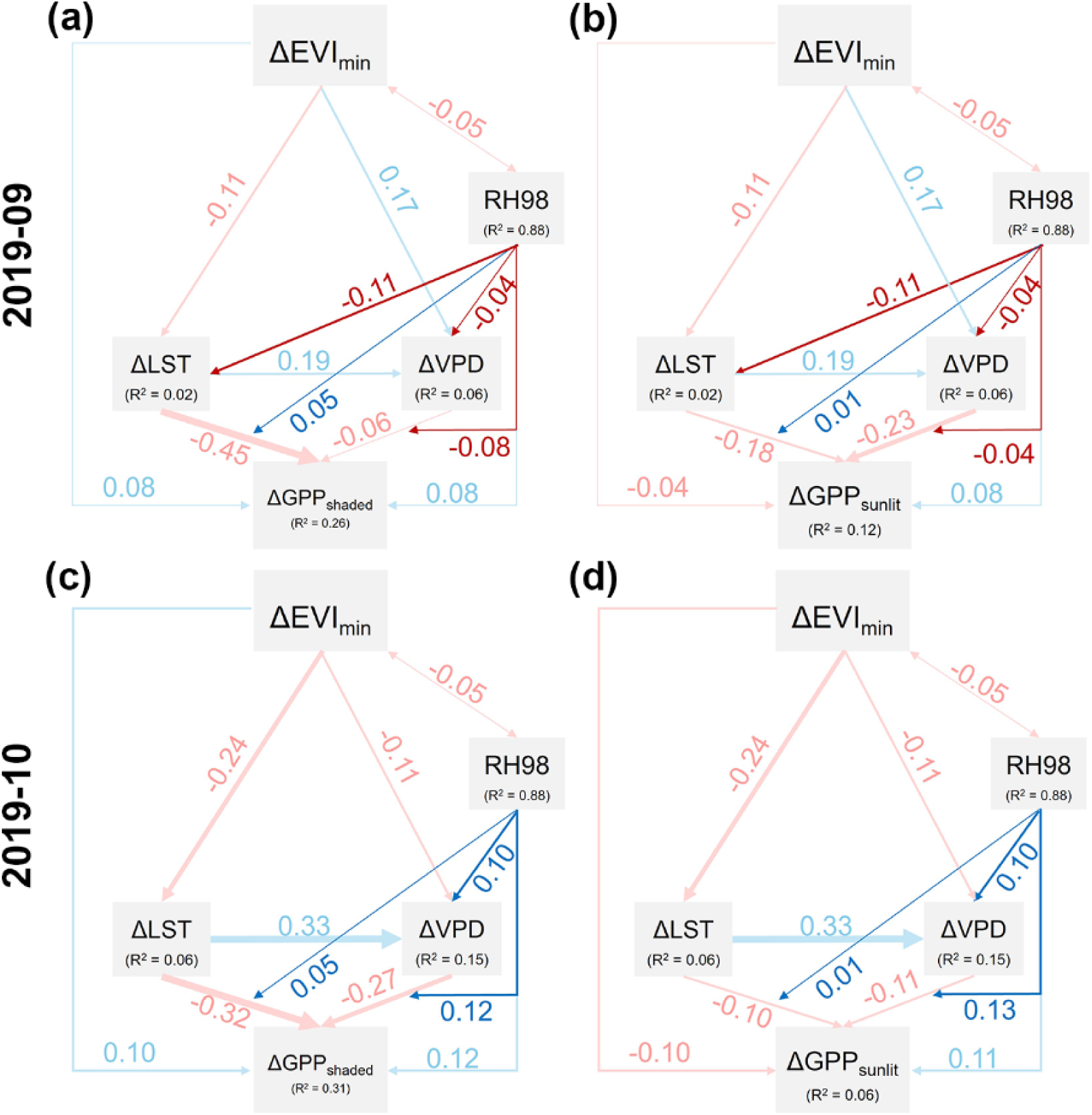
Piecewise Structural Equation Model (pSEM) illustrating the regulatory network for shaded and sunlit GPP anomalies. Each panel represents an independent pSEM with an identical theoretical structure, evaluated for: **(a)** shaded GPP anomaly and **(b)** sunlit GPP anomaly during the heatwave peak (Sept. 2019), and **(c)** shaded GPP anomaly and **(d)** sunlit GPP anomaly during the recovery phase (Oct. 2019). The models illustrate the complex interactions between hurricane legacy effects (ΔEVImin), forest structure (RH98), and environmental factors. Arrows represent significant causal paths (p≤0.05), with blue/light-blue and red/light-red indicating positive and negative effects, respectively; thinner lines denote weaker standardized path coefficients. Highlighted paths (red/blue arrows) emphasize the buffering and moderating effects of RH98.

Beyond the structural mediation of RH98, the pSEM also captured the direct impacts of hurricane disturbance intensity, represented by ΔEVImin, which is the minimum EVI anomaly within three months post-hurricane, used here as a proxy for immediate structural disturbance intensity (**Methods**, **Figure 2e, 3e, 4**). We observed a significant negative bi-directional interaction between ΔEVImin and RH98 (-0.05). Across all models, ΔEVImin exerted a consistent negative effect on ΔLST (-0.11 and -0.24 for September and October, respectively), suggesting that greater disturbance intensity led to increased surface warming. The influence of ΔEVImin on GPP exhibited contrasting directions between canopy layers. While more severe damage (more negative ΔEVImin) directly suppressed both total and shaded ΔGPP, it showed a negative path coefficient toward ΔGPPsunlit.

## Forest vulnerability to global hurricane-heatwave compound events

We extended our BSTS framework to 21 hurricane-heatwave events identified globally between 2004 and 2020 (**Methods)** to assess the global prevalence and determinants of enhanced forest vulnerability to the hurricane-heatwave compound events. Among these cases, significant negative hurricane legacy effects (where observed ΔGPP was significantly lower than the counterfactual prediction) were detected in 9 out of 21 events (42.9%, Hurricane Michael included). In the remaining 12 cases, the legacy effects during the subsequent heatwave were statistically non-significant. Notably, no cases exhibited positive legacy effects.

Categorizing the 21 global events by potential determinants identified three primary factors: the strength of heat and drought during the heatwave, and the soil type (**Figure 5**). The magnitude of the subsequent climate stress is a dominant driver. Events characterized by more extreme temperature anomalies and severe water deficits exhibited a substantially higher probability of manifesting negative hurricane legacy effects. Forests situated on loamy soils were found to be more susceptible to legacy effects. In contrast, forests on clayey or rocky soils appeared more resilient, showing a lower frequency of significant productivity declines during the post-hurricane heatwave. Compared to heatwave strength and soil type, hurricane intensity and the interval between disturbances did not provide additional information for prediction of heatwave vulnerability.

**Figure 5.**
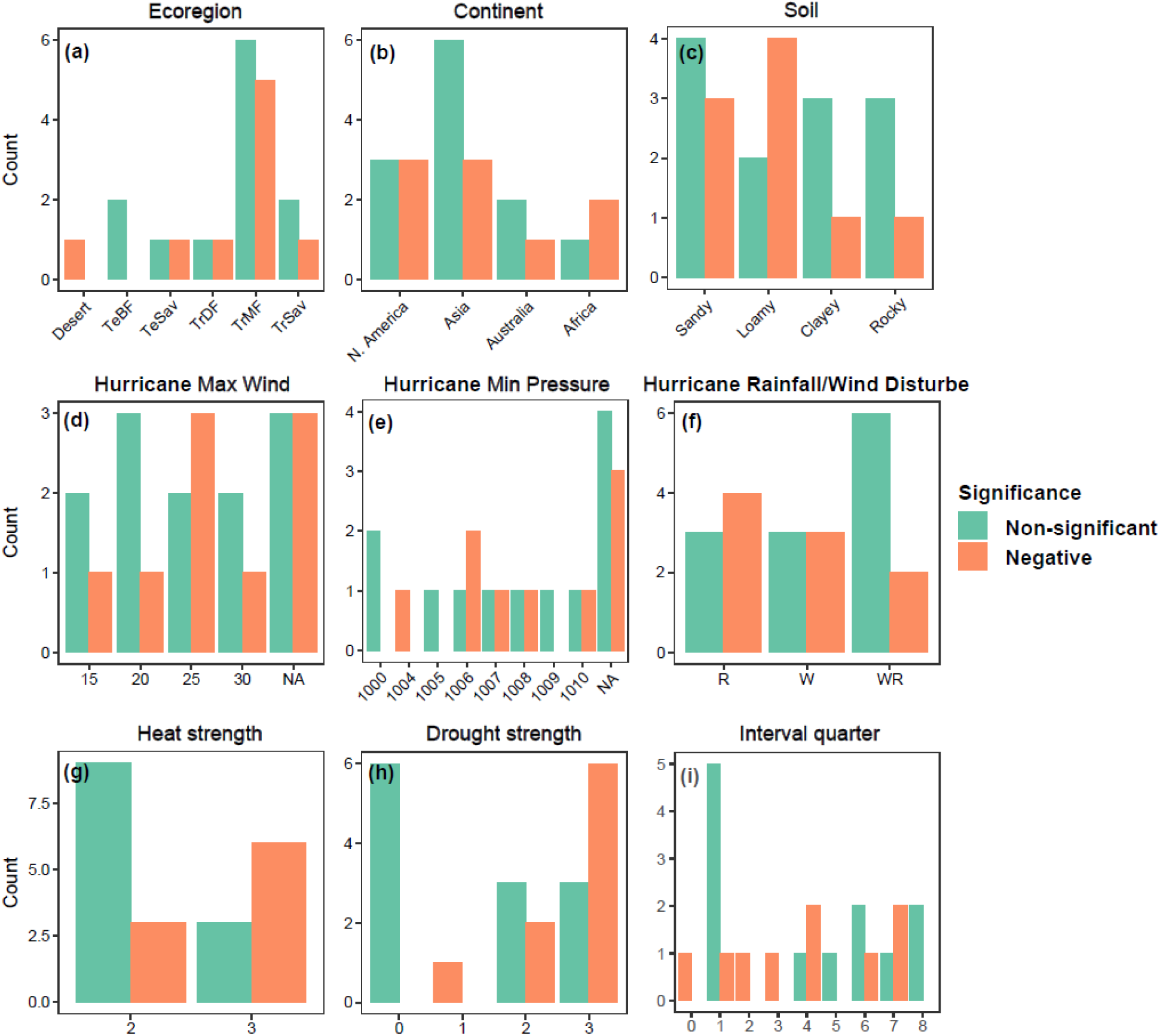
Global synthesis of ecological legacy effects across 21 hurricane-heatwave compound events. Bar charts illustrate the frequency of detecting significant negative hurricane legacy effects (orange) versus non-significant effects (green) across different categorical and discretized continuous variables. The events are stratified by ecological and geographical context: **(a)** Ecoregion, **(b)** Continent, and **(c)** Soil type; Hurricane Characteristics: **(d)** Maximum wind speed at landfall, **(e)** Minimum central pressure, and **(f)** Primary disturbance type, wind-dominant (W) vs. rainfall-dominant (R); Heatwave Severity: **(g)** Heat strength and Drought strength during the heatwave, and **(h)** the Interval time between hurricane landfall and heatwave.

## Discussion

Our study of Hurricane Michael confirms that prior hurricane disturbance significantly amplifies forest vulnerability to subsequent heatwaves. This legacy effect is largely mediated by the loss of forest vertical structure, i.e., the reductions in canopy height and structural diversity (**Figure 6**). Intact, tall canopies act as biological parasols, buffering sub-canopy microclimates against extreme temperature and water deficit anomalies^10, 11, 12, 20^. Meanwhile, high structural and component diversity facilitates diverse ecological niches and asynchrony in individual responses to climate fluctuation, thus maintaining forest stability^15, 16^. Furthermore, our results show that the shaded sub-canopy layers are more sensitive to temperature fluctuations than the sunlit leaves, presumably due to the lower light-saturation points and lower heat-tolerance of shaded leaves^14^. Most of the severe windthrow damage occurred in stands of ≥25 m height^8^. By removing the physical shelter, hurricanes leave these temperature-sensitive shaded leaves^21^ directly exposed to extreme stress.

**Figure 6.**
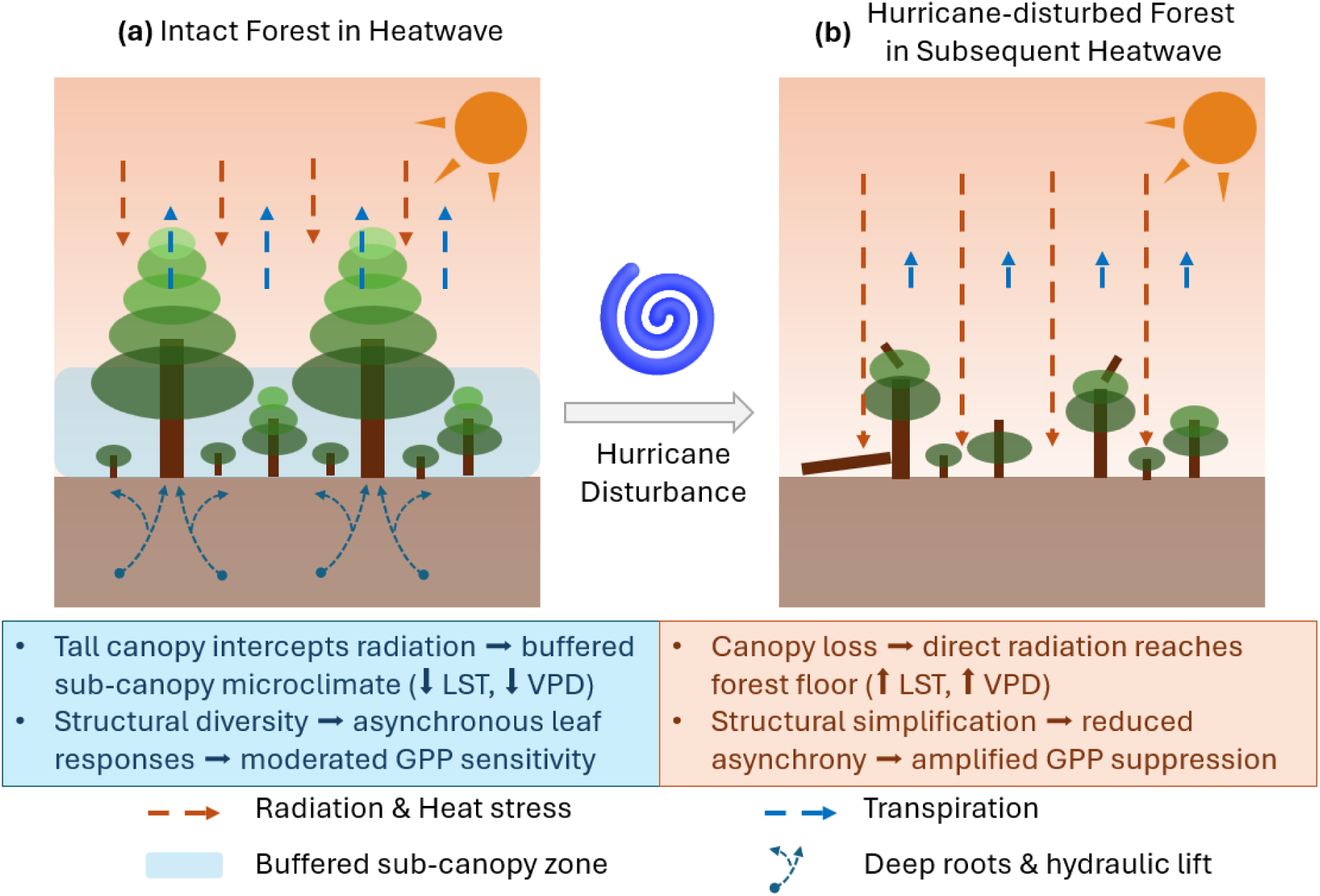
Conceptual schematic illustrating the mechanisms by which hurricane-induced structural simplification amplifies forest vulnerability to subsequent heatwaves. **(a)** An intact, tall-diverse forest intercepts incoming radiation at the canopy, buffering sub-canopy microclimate (reduced LST and VPD), while structural and species diversity promotes asynchronous leaf responses that moderate GPP sensitivity. Deep roots facilitate hydraulic lift, partially buffering soil water deficits; however, tall stature also imposes a hydraulic cost (longer transport pathways, higher transpiration demand) that can exacerbate VPD sensitivity under extreme heat. **(b)** Hurricane disturbance removes the canopy shelter, allowing direct radiation and heat stress to penetrate to the understory layer, exposing temperature-sensitive shaded leaves to extreme conditions. Structural simplification reduces buffering and ecological asynchrony, while disrupted root systems impair hydraulic regulation, collectively amplifying GPP suppression during the subsequent heatwave.

Interestingly, we found a reversal of the relationship between canopy height and forest VPD sensitivity between the heatwave peak and the recovery phase. We propose that this reversal is the result of a functional trade-off. On one hand, the shading effect of the tall canopies and the hydraulic lift function of deep roots^22, 23^ can enhance forests’ capacity to buffer hydrological stress. On the other hand, larger trees face greater hydraulic safety challenges due to longer water transport pathways and higher transpiration demands from larger leaf areas^24, 25, 26^. During the heatwave peak, extreme atmospheric demand likely pushes these tall-statured trees toward their hydraulic failure thresholds, where transport costs outweigh canopy buffering benefits, enlarging hydrological sensitivity. However, as conditions moderate during the recovery period, the regulatory advantages of complex structures regain dominance, effectively lowering the sensitivity. This highlights that forest structure is not a static buffer but a dynamic regulator whose effect is context-dependent^27^.

Our synthesis of 21 global hurricane-heatwave events suggests that potential structural damage from hurricanes is more likely to affect productivity when the ecosystem is pushed toward its coping capacity by intense heat and water stress. This explains why significant legacy effects are more frequently detected during high-intensity heatwaves. The lack of significant correlation with hurricane intensity or interval time in our study likely reflects a threshold effect within our sampling strategy: by selecting only hurricanes with R64 buffers (≥33 m/s), we focused on disturbances that are absolutely structurally destructive. In comparison, previous work identified potential cyclone disturbed vegetation with a threshold of 18 m/s^28^. Within a two-year window, these forests remain in the early stages of recovery, making the precise timing or variations in hurricane characteristics secondary to the severity of the subsequent heatwave. Furthermore, we found that forests on loamy soils are more prone to hurricane legacy effects than those on clayey or rocky soils, which may because of their acquisitive plant functional traits characterized by high productivity and transpiration demand^29, 30^. The acquisitive strategies can become a physiological liability once hurricane-induced structural damage impairs the hydraulic and microclimatic regulation required to meet those demands under stress^18^.

While our study provides robust evidence for structural mediation, some uncertainties remain. First, GEDI forest structure data that we used were only available post-Hurricane Michael. Although our pre-analysis confirmed that the sample and reference groups shared nearly identical trajectories of productivity and greenness for five years prior to the hurricane disturbance (greenness unshown), we cannot entirely rule out potential background variations like topography and pre-hurricane soil water storage that may affect the severity of hurricane disturbances^28, 31^. Second, the use of ΔEVImin as a proxy for hurricane damage intensity, while necessary due to the lack of high-resolution instantaneous wind field products, is a spectral manifestation of damage rather than a direct physical measure. Nonetheless, ΔEVImin serves as a robust integrative proxy, capturing the actual biological impact on the canopy that meteorological parameters alone might miss. Finally, the remote-sensing derived LST and VPD products primarily reflect the thermal and hydrological conditions at the canopy-top surface rather than the actual sub-canopy microclimate experienced by individual trees in the forests. During extreme heatwaves, true sub-canopy environments are typically characterized by lower temperatures and reduced water deficits compared to the exposed upper canopy, with the differences being more pronounced for more extreme climate anomalies^11^. Therefore, our reliance on satellite-derived temperatures may represent a conservative estimate of the buffering capacity of intact forests. These limitations likely account for the residual complexities in non-structural pathways within our pSEM, suggesting the influence of intermediate biogeophysical processes, such as nutrient cycling, that fall beyond the structural scope of this study. Future studies integrating in-situ microclimate sensors, understory LiDAR, or mechanistic microclimate modeling would further help unravel these hidden layers of the compound disturbance stories.

Compound disturbances can lead to impacts far exceeding the sum of individual stressors due to exceeded coping capacities and legacy effects^1, 5, 7^. Compound climate disturbances such as hurricane-heatwave events might have been considered rare "black swan" events in the past decades^32, 33^. However, growing observations and Earth System Models (ESMs) indicate that the frequency of such compound disturbances is increasing rapidly^32, 33, 34^. Our results suggest that hurricane-induced structural degradation acts as a critical mediator of forest vulnerability to subsequent heatwaves, creating a "1+1 > 2" compound effect on ecosystem productivity^3, 5^, particularly in loamy-soil regions facing increasingly intense extreme heatwaves in a warming climate. Ignoring these compound effects in ESMs could lead to a significant underestimation of the negative impacts of climate change on the global carbon cycle^3, 5, 7^. Effective forest management must transition from assessing single-driver risks to more integral frameworks accounting for non-linear responses to compound disturbances.

## Methods

### Data source and preprocessing

Hurricane information was obtained from the International Best Track Archive for Climate Stewardship (IBTrACS) project^35, 36^. IBTrACS is one of the most commonly used sources for tropical cyclone data, providing location, intensity, and size for all known tropical and subtropical cyclones at a resolution of 3 h, in which the R64 (the radius of 64-knot wind) and R50 (the radius of 50-knot wind) records are available after 2004^37^ (see Sampling Strategy). The monthly-scale GPP (gC m^-2^ month^-1^) data used in this study were obtained from the global Sunlit and Shaded GPP for vegetation canopies dataset^38^ with a resolution of 0.05°(∼5km) for the period from 1992 to 2020. LST data was obtained from MODIS MOD11A2.061 Terra land surface temperature and emissivity dataset^39^ with a resolution of 1km; VPD was obtained from TerraClimate Monthly Climate and Climatic Water Balance database with a resolution of ∼4.6km^40^. For Hurricane Michael only, the Enhanced Vegetation Index (EVI) was calculated with USGS Landsat 8 or Landsat 5 Collection 2 Level-2 tier following the equation reported by (Huete *et al.*, 2002)^41^. Forest canopy height was quantified as the relative height metrics at 98% (RH98) observed by the Global Ecosystem Dynamics Investigation (GEDI) mission L2A product, while forest structure diversity was represented by the foliar height diversity (FHD) in GEDI L2B product. Due to the limited reliability of GEDI RH98 retrievals for vegetation below 3 m, attributable to the LiDAR pulse width constraint of the sensor^42^, footprints with RH98 < 3 m were excluded from subsequent analyses.

All variables excluding the GEDI metrics were harmonized to a monthly temporal resolution and a 0.05° ×0.05° spatial grid, including GPP, VPD, LST and EVI. To isolate disturbance-induced signals from natural seasonal variability, we calculated a climatological baseline for each calendar month (January through December) for each pixel using the five-year period preceding hurricane (e.g., 2013.09 - 2018.09 for Hurricane Michael). Monthly anomalies were then computed as the difference between the observed value of each month and the corresponding long-term historical monthly mean value.

### Sampling strategy

We defined hurricane-disturbed forests as areas with woody cover exceeding 30% located within the mean R64 radius of each hurricane track point. Sustained wind speeds were at least 64 knots (∼33 m/s) within the R64 radius, this is considered a critical threshold for causing significant structural damage to both forest canopies and anthropogenic infrastructures^43^. In the hurricane track database, R64 values are recorded separately for four quadrants. For this study, we used the average R64 across available quadrants to simplify the spatial delineation of the disturbed zone while mitigating the impact of missing directional data in the historical records, ensuring a consistent and representative sampling area for each storm event.

An undisturbed reference group was set within a buffer zone of 10 to 50 nautical miles (nmile) beyond the R50 boundary. To ensure that the reference group was ecologically comparable to the disturbed group, we restricted selection to pixels with woody cover > 30% and an ecoregion within the two most dominant ecoregions within the sampling region. This strategy ensures that the reference group shares a similar environmental and ecological background with the disturbed forests while remaining sufficiently distant from the structural wind-damage region. This sampling strategy does not intend to capture the full spectrum of hurricane impacts but prioritizes the establishment of a statistically robust counterfactual to isolate the specific effects of hurricane disturbance on forest productivity.

### Statistical methods

To address Scientific Question 1, we employed a Bayesian Structural Time Series (BSTS) model to quantify the impact of Hurricane Michael on forest GPP^44, 45^. We utilized a five-year pre-disturbance period (2013.09 - 2018.09) to train the model, using time series of GPP, LST and VPD from the reference group as predictors. This model generated a counterfactual GPP scenario for the post-hurricane period (2018.10 - 2020.12), representing the expected forest productivity in the absence of the hurricane. By comparing this counterfactual baseline with the observed GPP, we identified significant negative deviations as "legacy effects". Specifically, we examined whether these legacy effects exacerbated the GPP decline during the subsequent extreme heatwave. The counterfactual approach allows us to quantify vegetation disturbances and recovery under an unstable background^44^, thus is exactly suitable for our study assessing forest performance during hurricane-heatwave compound events.

To address Scientific Question 2, we first quantified structural changes. Since GEDI data only became available in March 2019 (five months after hurricane landfall), we characterized hurricane-induced structural disturbance by calculating the divergence in canopy height (RH98) and Foliage Height Diversity (FHD) between the Hurricane-disturbed group and the reference group. We performed linear regressions to quantify the relationship between RH98 and anomalies in LST (ΔLST) and VPD (ΔVPD) separately during the peak heatwave (2019.09) and the subsequent recovery phase (2019.10). To assess the sensitivity of forest productivity to environmental stress, we used grouped regression models. All pixels were binned into 10-meter intervals based on their RH98, and the slopes of ΔGPP versus ΔLST and ΔVPD were calculated for each bin to quantify how height-dependent structure modulates sensitivity to heat stress.

Finally, we constructed a piecewise Structural Equation Model (pSEM)^46^ for the sample region only to integrate the complex pathways between hurricane disturbance, forest structure, thermal-hydrological stress, and forest productivity (total, shaded, and sunlit GPP). Path coefficients in the pSEM are standardized, representing the expected change in the response variable in standard deviation units associated with a one standard deviation change in the predictor, thereby allowing direct comparison of effect sizes across paths. In the pSEM, the minimum EVI anomaly (ΔEVImin) within three months post-hurricane was used as a proxy for immediate hurricane disturbance intensity to reduce the impact of potential temporal lags in remote sensing detection. Due to the high collinearity between RH98 and FHD (Pearson correlation coefficient: 0.74), only RH98 was retained to represent forest structure. The priori model assumed a Bi-directional interaction between ΔEVImin and RH98, reflecting both the higher susceptibility of tall forests to windthrow^31, 47, 48^ and the reduction of canopy height following disturbance^8, 48^; direct impacts of both ΔEVImin and RH98 on ΔLST, ΔVPD, and ΔGPP, reflecting potential legacy effect of hurricane through forest structure and other paths, such as forest components and plant functional traits^47, 49^; Causal paths where ΔLST influences ΔVPD, and both jointly affect ΔGPP. We are specifically interested in the buffering effect and the moderating effect of forest structure, i.e., the expected negative impacts of RH98 on ΔLST and ΔVPD, and RH98’s role in altering ΔGPP sensitivity to ΔLST/ΔVPD via interaction terms during heatwave.

To address Scientific Question 3, we screened all hurricane-heatwave compound events occurring within a 24-month interval between 2004 and 2020 at global scale. Hurricanes were identified where the R64 buffer of a track point intersected terrestrial forests; heatwaves were defined as months where the temperature anomaly exceeded two standard deviations of the historical monthly mean. Out of 50 identified compound events, 29 were excluded due to an insufficient area of suitable control group, resulting in 21 cases for detailed analysis. For each event, we applied the BSTS model similar to Scientific Question 1 to detect significant legacy effects of hurricanes. We subsequently evaluated how factors such as hurricane intensity, heatwave magnitude, interval time, vegetation type, and soil characteristics influenced the probability of detecting a significant hurricane legacy effect.

All data acquisition and initial spatial-temporal aggregation tasks were performed on the Google Earth Engine platform (Google, Mountain View, CA, USA), while subsequent data preprocessing, statistical analyses, and visualization were conducted using R 4.5.1 (R Foundation for Statistical Computing). The BSTS models were implemented using the “CausalImpact” package, the pSEM model was implemented with the “piecewiseSEM” package, and all the data visualization were generated using the “ggplot2” package.

## Author contributions

J.D. conceived the study, designed the methodology, performed the analyses, and wrote the manuscript. A.B.H. supervised the project and contributed to study design and manuscript revision. X.L. contributed to data processing. G.J.K., M.U., and T.L.M. contributed to interpretation of results and manuscript revision. All authors reviewed and approved the final manuscript.

## Acknowledgments

The authors acknowledge support from the U.S. Department of Energy, Office of Science, Office of Biological and Environmental Research, under Award Number DE-SC0025272.

## Data availability

The data used in this study are all publicly available from their original sources. GPP data are available at https://doi.org/10.5061/dryad.dfn2z352k; LST data at https://doi.org/10.5067/MODIS/MOD11A2.061; VPD data from TerraClimate at https://www.climatologylab.org/terraclimate.html; hurricane track data (IBTrACS v4.01) at https://doi.org/10.25921/82ty-9e16; GEDI L2A and L2B products at https://doi.org/10.5067/GEDI/GEDI02_A.002 and https://doi.org/10.5067/GEDI/GEDI02_B.002, respectively. No new data products were generated in this study.

## Code availability

The code used in this study is available at https://doi.org/10.5281/zenodo.20148725 and will be made publicly accessible upon acceptance of the manuscript.

